# Plasmid-free cheater cells commonly evolve during laboratory growth

**DOI:** 10.1101/2023.05.19.541508

**Authors:** Amber M. Bedore, Christopher M. Waters

**Affiliations:** Department of Microbiology and Molecular Genetics Michigan State University, East Lansing, Michigan, USA, 48824

**Author notes:** Corresponding Author: 5180 Biomedical and Physical Sciences 567 Wilson Road East Lansing, MI 48824.

**Keywords:** Beta-lactamase, aminoglycoside phosphotransferase, efflux pumps, plasmid maintenance, public goods

## Abstract

It has been nearly a century since the isolation and use of penicillin, heralding the discovery of a wide range of different antibiotics. In addition to clinical applications, such antibiotics have been essential laboratory tools, allowing for selection and maintenance of laboratory plasmids that encode cognate resistance genes. However, antibiotic resistance mechanisms can additionally function as public goods. For example, secretion of beta-lactamase from resistant cells, and subsequent degradation of nearby penicillin and related antibiotics, allows neighboring plasmid-free susceptible bacteria to survive antibiotic treatment. How such cooperative mechanisms impact selection of plasmids during experiments in laboratory conditions is poorly understood. Here, we show that the use of plasmid-encoded beta-lactamases leads to significant curing of plasmids in surface grown bacteria. Furthermore, such curing was also evident for aminoglycoside phosphotransferase and tetracycline antiporter resistance mechanisms. Alternatively, antibiotic selection in liquid growth led to more robust plasmid maintenance, although plasmid loss still occurred. The net outcome of such plasmid loss is the generation of a heterogenous population of plasmid-containing and plasmid-free cells, leading to experimental confounds that are not widely appreciated.

**Importance:** Plasmids are routinely used in microbiology as readouts of cell biology or tools to manipulate cell function. Central to these studies is the assumption that all cells in an experiment contain the plasmid. Plasmid maintenance in a host cell typically depends on a plasmid-encoded antibiotic resistance marker, which provides a selective advantage when the plasmid containing cell is grown in the presence of antibiotic. Here we find that growth of plasmid-containing bacteria during laboratory conditions in the presence of three distinct antibiotic families leads to the evolution of a significant number of plasmid-free cells, which rely on the resistance mechanisms of the plasmid-containing cells for viability. This process generates a heterogenous population of plasmid-free and plasmid-containing bacteria, an outcome which could confound further experimentation.

## Introduction

Cooperation among and between bacterial species is a key element for survival. Members of bacterial communities, whether in single or mixed populations, communicate and coordinate metabolism and other cellular processes to survive a changing environment. This cooperative social dynamic is well characterized in bacteria, and examples include bacterial biofilms, which protect a population from environmental insults by the cooperative formation of an extracellular matrix, and quorum sensing, which involves cell communication to induce cooperative behaviors (1–3).

An important aspect of bacterial cooperation is the production and distribution of shared extracellular metabolites, nutrients, and proteins. Collectively referred to as public goods, these products are available to and benefit all members of the population, either with or without reciprocal behavior from non-producing recipients. The production of enzymes to break down extracellular nutrients, secretion of siderophores for iron acquisition, and biosurfactants for bacterial motility are all examples of public goods that bacteria produce to benefit themselves and others in the population (4–6).

Public goods represent a cooperative behavior in bacteria as they are costly to produce but do not require reciprocation from recipients. Based on these traits, members that produce and distribute public goods are considered cooperators, and those who benefit from the cooperative behavior, but do not contribute to the population, are referred to as cheaters (7, 8). Social cheating is well documented in bacteria, and cooperative members have even developed molecular mechanisms such as kin-recognition and policing to prevent cheaters from persisting in the population (9, 10). Antibiotic resistance mechanisms can also function as public goods, an example being secreted beta-lactamases.

Beta-lactamase, an enzyme encoded by the *bla* gene, is utilized by bacteria to cleave the beta-lactam ring of penicillin-class antibiotics (11, 12). This enzyme is secreted by resistant bacterial cells, resulting in degradation of environmental penicillins. This detoxification allows susceptible members of the community to grow (13–17). In this way, beta-lactamases can be considered public goods as they are manufactured by cooperative cells that secrete them into the environment to destroy antibiotics that are inhibiting the growth of susceptible members of the community. But the evolution of social cheaters for this public good in laboratory experiments that utilize *bla* and beta-lactams for plasmid selection has not been reported.

Natural plasmids have several mechanisms to ensure they are not cured from bacterial populations including toxin/antitoxin and partitioning systems (18–21). However, to reduce their size for ease of manipulation, most laboratory plasmids lack such mechanisms and are rather maintained by encoding antibiotic resistance genes and growth of plasmid-containing bacteria in the appropriate antibiotic. The assumption is that any cells cured of the plasmid will be killed by the antibiotic. As plasmids are used for a variety of molecular biology processes such as gene reporters, overexpression constructs, protein purification, and genetic complementation, it is imperative for proper experimental interpretation that all members of the population under study encode the plasmids.

We have recently employed a cyclic di-GMP (c-di-GMP) biosensor in *Vibrio cholerae* in which translation of TurboRFP is enhanced by c-di-GMP binding to an RNA riboswitch (22, 23). This biosensor is located on a plasmid containing *bla* as an antibiotic marker to allow selection for plasmid maintenance. During studies of *V. cholerae* encoding this plasmid, we observed both fluorescent and non-fluorescent cells when the bacteria were grown as colonies on solid agar. Further investigation revealed the non-fluorescent cells were a population of cheaters that had lost the plasmid and exploited the detoxification of ampicillin in the growth media produced by beta-lactamase secretion by cooperative members of the population that maintained the plasmid. Subsequent analysis found that two different resistant mechanisms, an aminoglycoside phosphotransferase (24, 25) and a tetracycline antiporter (26), also demonstrated plasmid loss when bacteria were grown on a surface and, to a lesser extent, in liquid culture. Together, our results demonstrate antibiotic selection for plasmid maintenance during laboratory experimentation, especially when using surface grown bacteria, can lead to plasmid-free cheater cells, creating a heterogenous bacterial population.

## Results

### A c-di-GMP biosensor reveals a heterogenous population of cells during ampicillin selection

Biosensors are useful tools for measuring cellular metabolites at the level of single cells. An RNA-based biosensor, P*be_amcyan_Bc3-4_turborfp*, for the second messenger cyclic di-GMP, which is a global signal that controls biofilm formation and motility in bacteria (27), was recently developed in which expression of TurboRFP is controlled by two c-di-GMP binding tandem riboswitches (23). For this biosensor, TurboRFP production positively correlates with intracellular c-di-GMP concentrations (23). During studies using the P*be_amcyan_Bc3-4_turborfp* biosensor encoded on the plasmid pRP0122 (hereafter referred to as pRP0122) to quantify c-di-GMP levels in two *Vibrio cholerae* strains, the wild type (WT) strain and a Δ*vpsL* biofilm deficient mutant, we observed that large “spot” colonies exhibited a distinct color pattern. These spot colonies were formed by plating 10 μL of culture onto an agar plate containing ampicillin (100 μg/mL) to ensure plasmid maintenance. The colonies had a bright red center, surrounded by a white ring of cells with red bursts of cells dispersed throughout the outer white ring (Fig. 1). Such color patterns of colony morphology were generally consistent across both strains. As increased TurboRFP production indicates elevated c-di-GMP concentrations, we hypothesized that *V. cholerae* cells growing in a colony exhibited heterogeneity of c-di-GMP production.

**Fig. 1.**
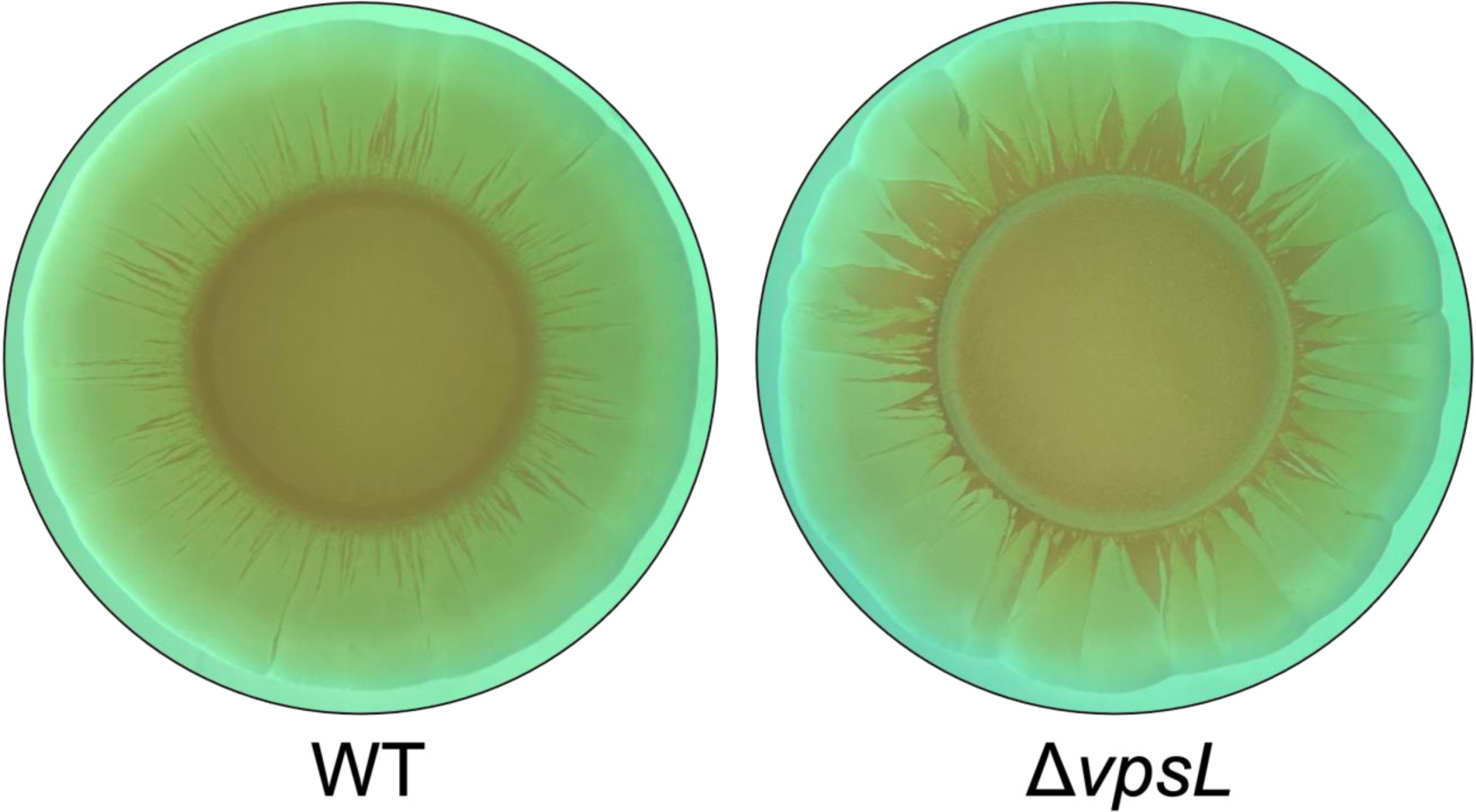
Ten μL of *V. cholerae* WT or Δ*vpsL* strains harboring the pRP0122 biosensor-encoding plasmid were spotted onto an LB plate containing 100 μg/mL ampicillin and grown for 16 hours at 35 °C. Non-fluorescent imaging of the colony showed a heterogenous phenotype of TurboRFP expression. Representative colonies are shown.

### Cells at the white edge of the colony have similar concentrations of c-di-GMP compared to cells at red center

To test if cells in the center of the colony had elevated c-di-GMP, cells from the red center and white ring of the colony were collected with a sterile loop and resuspended in LB broth. The resuspended cells were normalized to an OD_600_ of 0.1. Nucleotides were extracted and the concentration of c-di-GMP was determined using liquid chromatography tandem mass spectrometry (LC-MS/MS). However, our hypothesis was not supported as cells isolated from the white edge had comparable c-di-GMP concentrations to cells from the red center of the colony in both the WT and Δ*vpsL* strains (Fig. 2). Importantly, all samples had high concentrations of c-di-GMP that were greater than 5 μM, which should be sufficient to activate TurboRFP expression from the pRP0122 c-di-GMP biosensor.

**Fig. 2.**
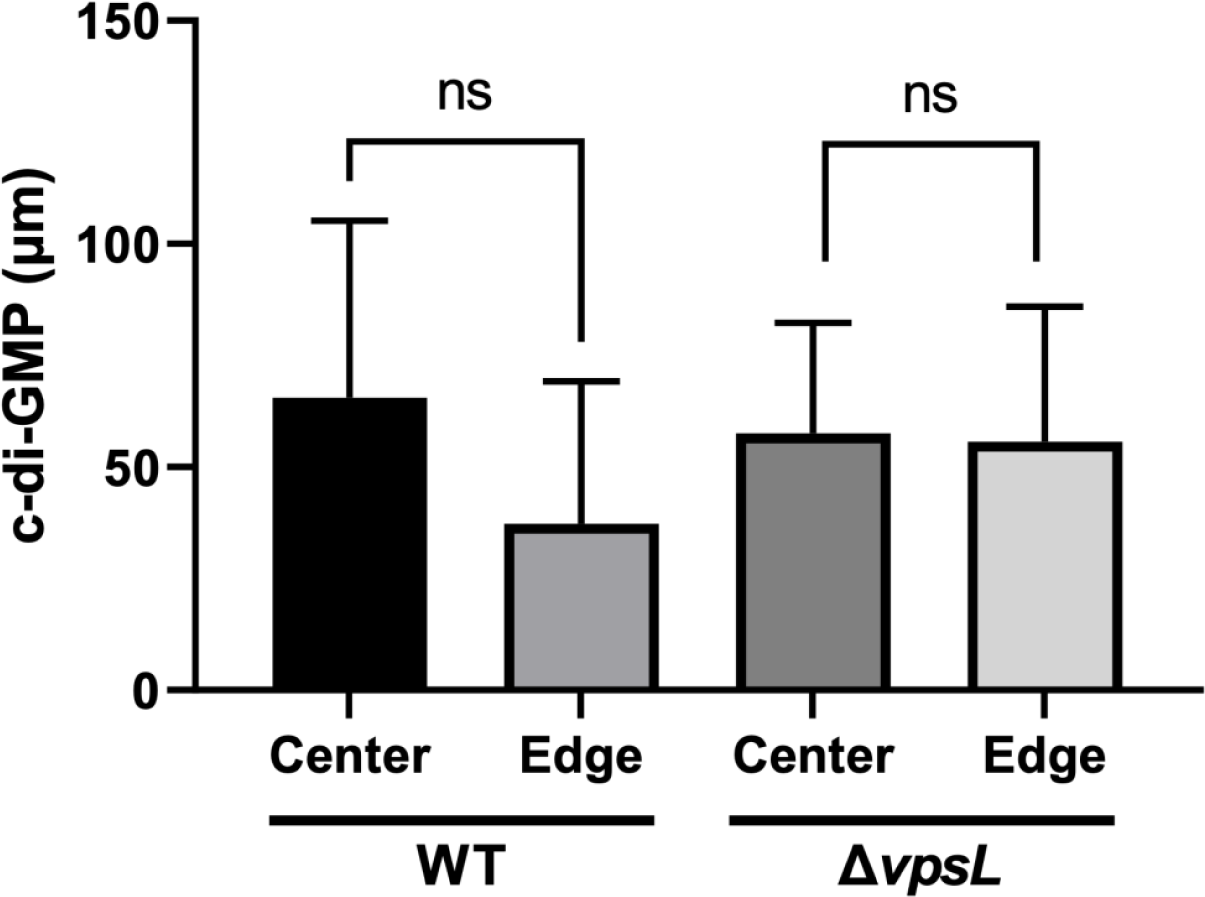
C-di-GMP was quantified using LC-MS/MS from biosensor containing cells collected from either the center or edge of a WT or Δ*vpsL* spot colony as shown in Fig. 1. Statistical significance was determined using a Student’s t-test (n=3), **ns** = not significant.

### Colony heterogeneity is due to plasmid loss

Given the similarities in c-di-GMP concentrations, we next hypothesized that the different colony morphologies were due to loss of the plasmid-encoded c-di-GMP biosensor. pRP0122 encodes the *bla* ampicillin resistance gene, and in previous experiments plasmid containing cells were selected using 100 μg/mL ampicillin. Considering that beta-lactamase can be a public good (13-15), we predicted that secreted beta-lactamases from the colony center degraded the antibiotic in the media allowing for the rise of plasmid-free cheater cells at the edge of the colony. Cooperative cells then assume the costs of carrying the plasmid, allowing plasmid free cells to dominate growth on the edge of the colony.

To begin to test this model, *V. cholerae* Δ*vpsL* colonies were propagated on a gradient of antibiotic concentrations to increase the selective pressure for plasmid maintenance. If the occurrence of cheaters is due to antibiotic degradation by beta-lactamases, then the frequency of cheaters would be inversely related to the concentration of antibiotic. When colonies were incubated on ampicillin, a large white perimeter of cheater cells grew around central cooperative red cells at and below the standard laboratory working concentration of 100 µg/mL that is commonly used for plasmid selection and maintenance (Fig. 3). The incidence of cheaters as evidenced by a smaller white perimeter decreased at concentrations 2- and 5-times the working concentration, though the white ring of cheater cells was never fully eliminated at any concentration of ampicillin.

**Fig. 3.**
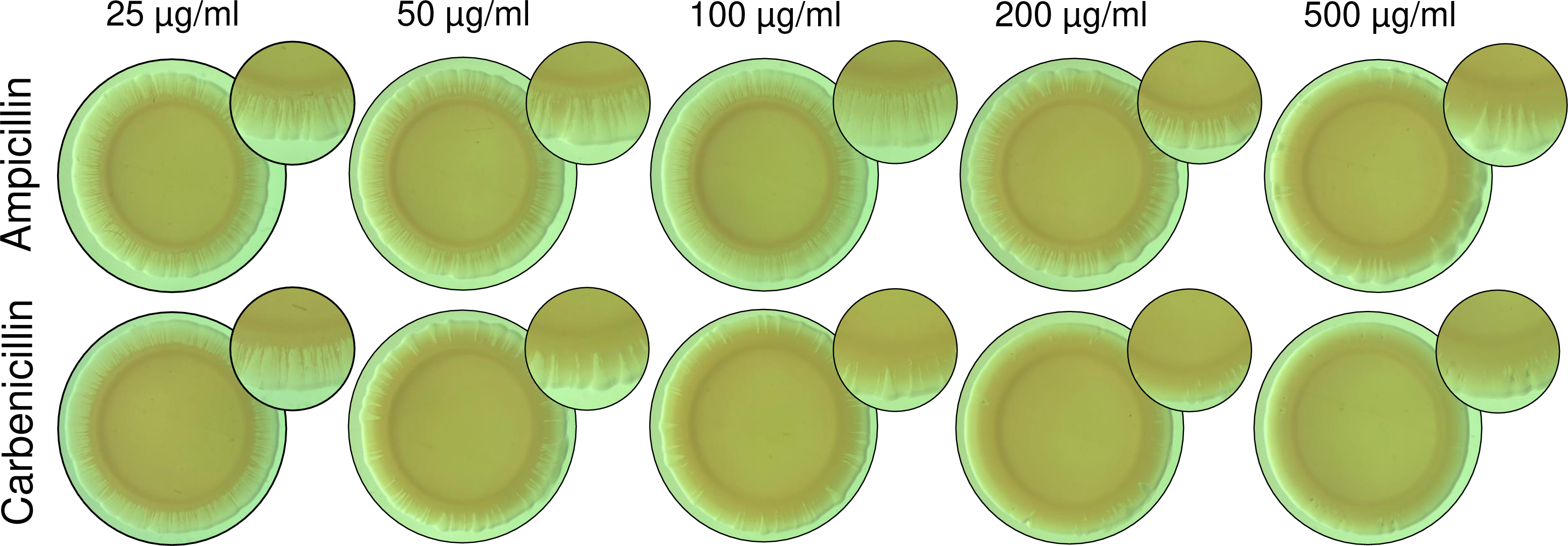
*V. cholerae* Δ*vpsL* with the plasmid c-di-GMP biosensor (pRP0122) was propagated on a gradient of increasing concentrations of either ampicillin or carbenicillin. A representative colony is shown, and the inlet provides a closer look at the edge of the colony.

We further tested this model by selecting for plasmid maintenance with carbenicillin, which is a widely used alternative to ampicillin due to its increased temperature and acid stability, and greater resistance to beta-lactamase degradation (28). Carbenicillin showed a decreased incidence of cheaters at the standard laboratory working condition of 100 µg/mL compared to ampicillin. When colonies were grown on carbenicillin at 2- and 5-times the working concentration, cheater cells were much less prominent than at these concentrations of ampicillin, as shown by the reduction of the white perimeter of cheaters surrounding cooperative red cells.

Importantly, at the working concentration of carbenicillin, cheater incidence is significantly reduced; however, a comparable frequency of cooperative cells does not appear until twice the working concentration of ampicillin. These results demonstrate that carbenicillin, the more stable antibiotic, inhibits the rise of cheater cells. Interestingly, cheaters were never fully eliminated in any conditions and are still present even at the highest concentration of carbenicillin. This suggests that the beta-lactamase enzymes secreted from this plasmid can degrade 5-times the working concentration of both of these penicillin-class antibiotics to permit the growth of plasmid-free cheaters.

### Single-cell analysis shows plasmid loss occurring in both the center and edge of spot colonies

The above experiments, which relied on colony visualization, qualitatively suggested that selection with ampicillin or carbenicillin at the standard 100 µg/mL working concentration led to plasmid loss. To quantify plasmid loss across the colony at an individual cell level, we performed single-cell quantification of TurboRFP from samples collected from the white edge and red center of WT and Δ*vpsL* colonies. Fluorescent and brightfield images were used in tandem to measure the number of plasmid-free cheater cells present that did not fluoresce with RFP, suggesting a loss of plasmid had occurred (Supplemental Fig. 1). Only 48.4% and 39.4% of the cells from the red center of the WT and Δ*vpsL* colony, respectively, exhibited RFP fluorescence, indicating they maintained the plasmid (Fig. 4). Therefore, in both cases the non-fluorescing cheater cells were a majority of the populations. Over 98.0% of cells from the white edge for both strains were non-fluorescent, suggesting that this region of the colony was dominated by plasmid-free cheater cells. When the entire colony was assessed, the frequency of non-fluorescent cells was similar to those collected from the red center, and again the non-fluorescent cheater cells were the majority of the population (Fig. 4).

**Fig. 4.**
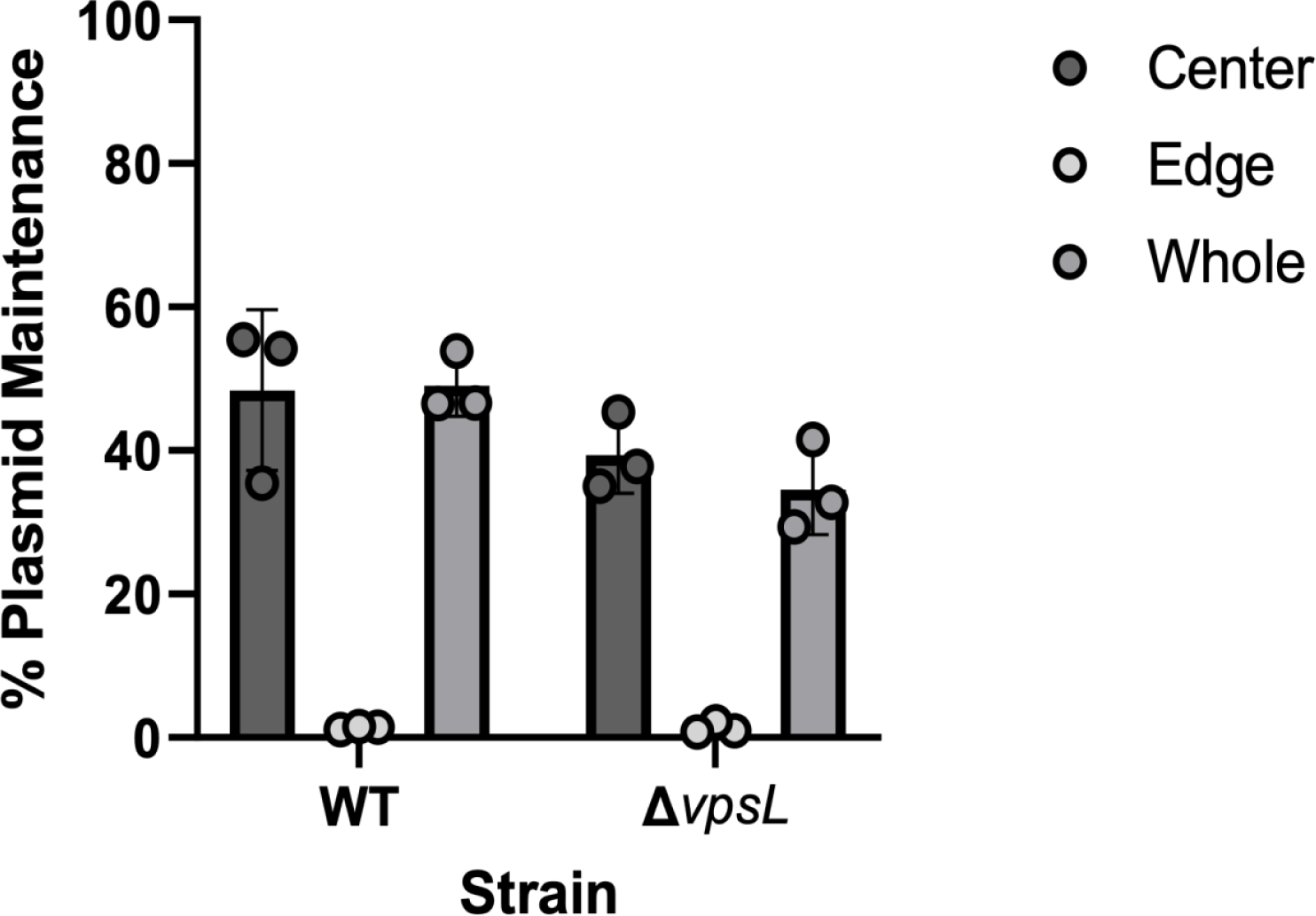
Single cells isolated from the center, edge, and the entire mixed colony were visualized for fluorescence. Non-fluorescent cells represent potential plasmid-free cheaters. The frequency of plasmid retention was calculated by dividing the total amount of fluorescent cells by the total amount of cells in brightfield and multiplying by 100 (n=3).

Taken together, these results support that cells growing at the center of the colony are cooperative cells that are carrying the cost of producing and secreting beta-lactamase that degrades the antibiotic in the growth media. However, a significant number of cells in the center, and more predominantly at the edge of the colony, are plasmid-free cheater cells that exploit the detoxification of the environment by cooperators and dominate growth at the edge of the colony.

### Plasmid loss occurs for selection of multiple antibiotics on multiple plasmid backbones

To test if plasmid loss only occurs upon selection utilizing a beta-lactamase resistance mechanism, we explored plasmid maintenance using a kanamycin *aph(3’)-I* resistance gene. *aph(3’)-I*, which encodes an aminoglycoside phosphotransferase, functions by phosphorylating an aminoglycoside to inactivate it, and this enzyme is located in the cell cytoplasm and not secreted from the cell (29, 30). We therefore hypothesized that selection for kanamycin resistance to maintain the plasmid would be more efficient than what we observed for selection with *bla* and ampicillin.

To directly compare these two resistance mechanisms, we replaced an *aph(3’)-I* resistance gene on a second plasmid (pEVS143) that produces GFP with a *bla* resistance gene. Importantly, pEVS143 is a different plasmid backbone than pRP0122 and encodes a p15A medium copy *ori* versus the RSF1010 low copy *ori* of pRP0122. Total cells harboring pEVS143 (encoding *aph(3’)-I* and selected with 100 μg/mL kanamycin) were analyzed from a whole colony using brightfield and fluorescence microscopy. Unexpectedly, we found that, when comparing total cells to fluorescent cells, only 73.6% of WT and 53.9% of Δ*vpsL* cells maintained the plasmid (Fig. 5A). When the antibiotic marker was changed to *bla* (pEVS143 [*bla*]), plasmid maintenance frequency dropped to 59.1% and 49.6% for WT and Δ*vpsL*, respectively. These results suggest that phosphorylation of kanamycin by the *aph(3’)-I* resistance gene decreased the overall antibiotic available for selection, leading to an intracellular-based mechanism of public goods that selected for plasmid loss when growing on a surface. These results also suggest that plasmid loss can be observed in two different plasmid backbones.

**Fig. 5.**
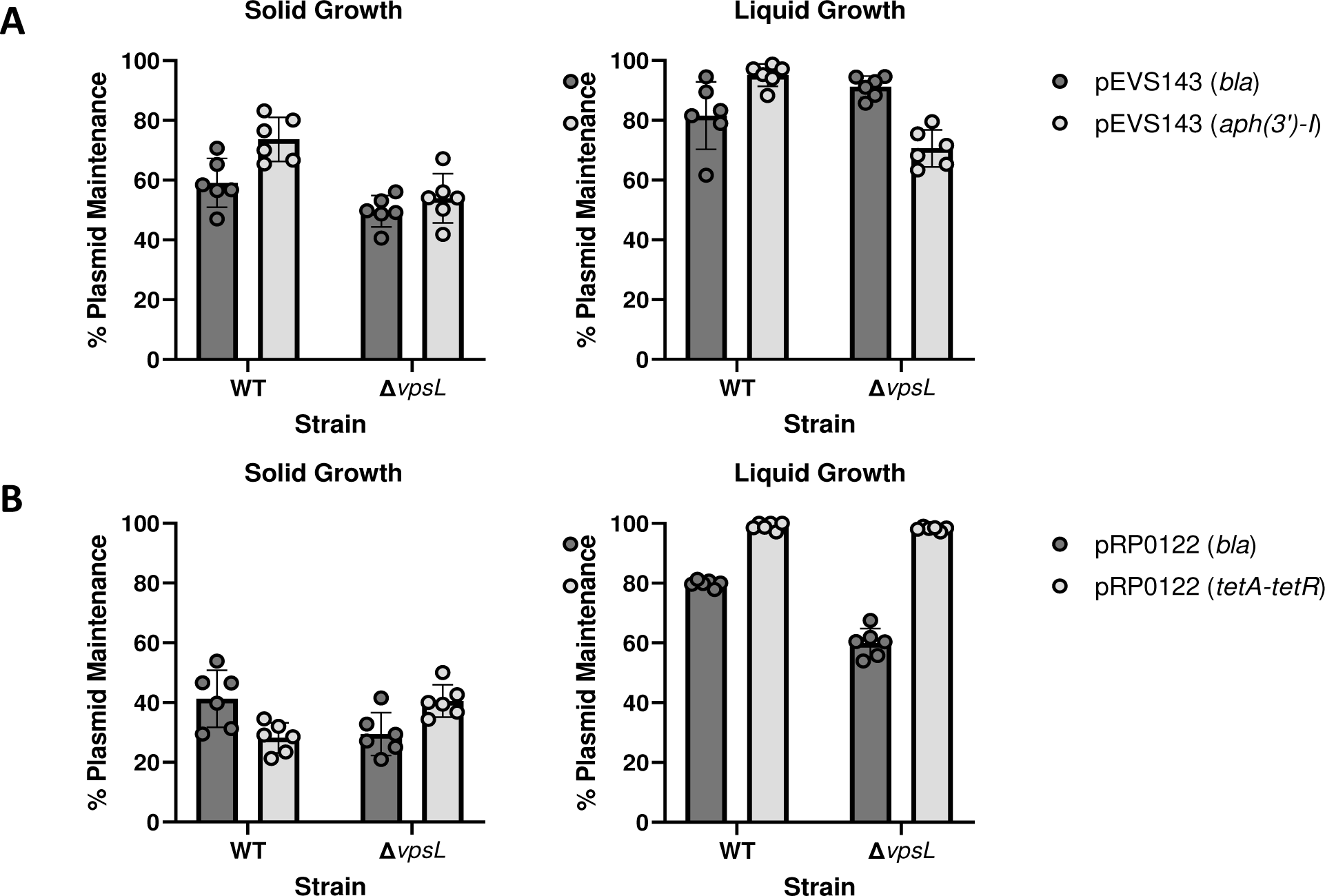
Plasmid loss was determined by growing either pEVS143 (**A**) or pRP0122 (**B**) strains with *bla*, *aph(3’)-I*, or *tetA-tetR* markers on solid (**left**) or in liquid (**right**) media and visualizing fluorescence. Non-fluorescent cells represent potential plasmid-free cheaters. The frequency of plasmid retention was calculated by dividing the total amount of fluorescent cells by the total amount of cells in brightfield and multiplying by 100 (n=3).

We further explored the generality of these results by performing similar experiments with the *tetA* resistance gene that does not enzymatically change the overall concentration of antibiotic. Rather, TetA is an efflux pump that removes tetracycline from inside the cell. We replaced the *bla* gene in the c-di-GMP biosensor plasmid pRP0122 with the *tetA-tetR* efflux pump system (pRP0122 [*tetA-tetR*]) to measure the effect of plasmid loss when the antibiotic is not degraded or inactivated. TetR represses *tetA* expression, and repression is relieved when antibiotic binds the TetR repressor to allow expression of *tetA*. When comparing fluorescent cells to total cells, we found that the efflux pump resistance mechanism of *tetA-tetR* led to only 28.2% plasmid maintenance in WT and 40.5% in Δ*vpsL* (Fig. 5B.). These results demonstrated that selection for plasmid maintenance with all three of these antibiotic resistance mechanisms was insufficient to completely eliminate the emergence of plasmid-free cheater cells when the bacteria were grown on solid media.

We next wondered if plasmid loss in liquid media using beta-lactam degradation, aminoglycoside inactivation, or efflux pumps led to the rise of cheaters in a planktonically grown cultures. To test this, we measured the frequency of plasmid maintenance first in the pEVS143 backbone encoding either *aph(3’)-I* or *bla* when grown overnight in liquid media with shaking in the presence of their respective antibiotics. In liquid growth, plasmid maintenance was increased to a range of 72.0-95.1% for both ampicillin and kanamycin selection, although kanamycin was more effective to maintain plasmids for the WT strain while ampicillin was more effective for the Δ*vpsL* strain (Fig. 5A).

We performed a similar analysis with the pRP0122 plasmid backbone, and again observed enhanced plasmid maintenance in liquid versus surface growth (Fig. 5B). Although significant plasmid loss was observed for ampicillin selection with plasmid maintenance at 79.8% and 59.9% for the WT and Δ*vpsL* strains, respectively, we found that selection with *tetA* significantly reduced the percentage of cheaters, with plasmid maintenance measured at 95.1% (WT) and 99.0% (Δ*vpsL*) (Fig. 5B). Nevertheless, even though selection for plasmid loss is greatly reduced in liquid cultures, plasmid loss still occurs both planktonically and on solid media.

### Loss of fluorescence correlates with loss of antibiotic resistance

Our previous results estimated plasmid loss through quantifying non-fluorescent cells. However, fluorescence could be lost either due to mutation or decreased expression of the fluorescence gene while the plasmid itself was still maintained. To confirm that the loss of fluorescence was an accurate readout of plasmid loss, we calculated the frequency of plasmid loss using another genetic marker on the plasmid, antibiotic resistance. Whole colonies were resuspended and plated onto non-selective media and allowed to grow into isolated colonies. These colonies were then patched onto non-selective and selective media and analyzed for growth.

When colonies were plated on non-selective media, two distinct colony morphologies were observed in the populations containing the pRP0122 (*bla*) and pRP0122 *(tetA-tetR*) c-di-GMP containing plasmids. Small colonies of pRP0122 (*bla*) and pRP0122 (*tetA-*tetR) were visibly red, while larger colonies were white (Fig. 6). This result suggests maintaining the c-di-GMP biosensor exerts a significant cost to the cell. A similar phenomenon of green and white colonies was observed in strains containing either the pEVS143 (*aph(3’)-I*) or pEVS143 (*bla*) plasmid, but the colonies were similar in size, suggesting pEVS143 (*aph(3’)-*I) does not impair as high of a fitness cost as pRP0122 (*bla*) (Fig. 6).

**Fig. 6.**
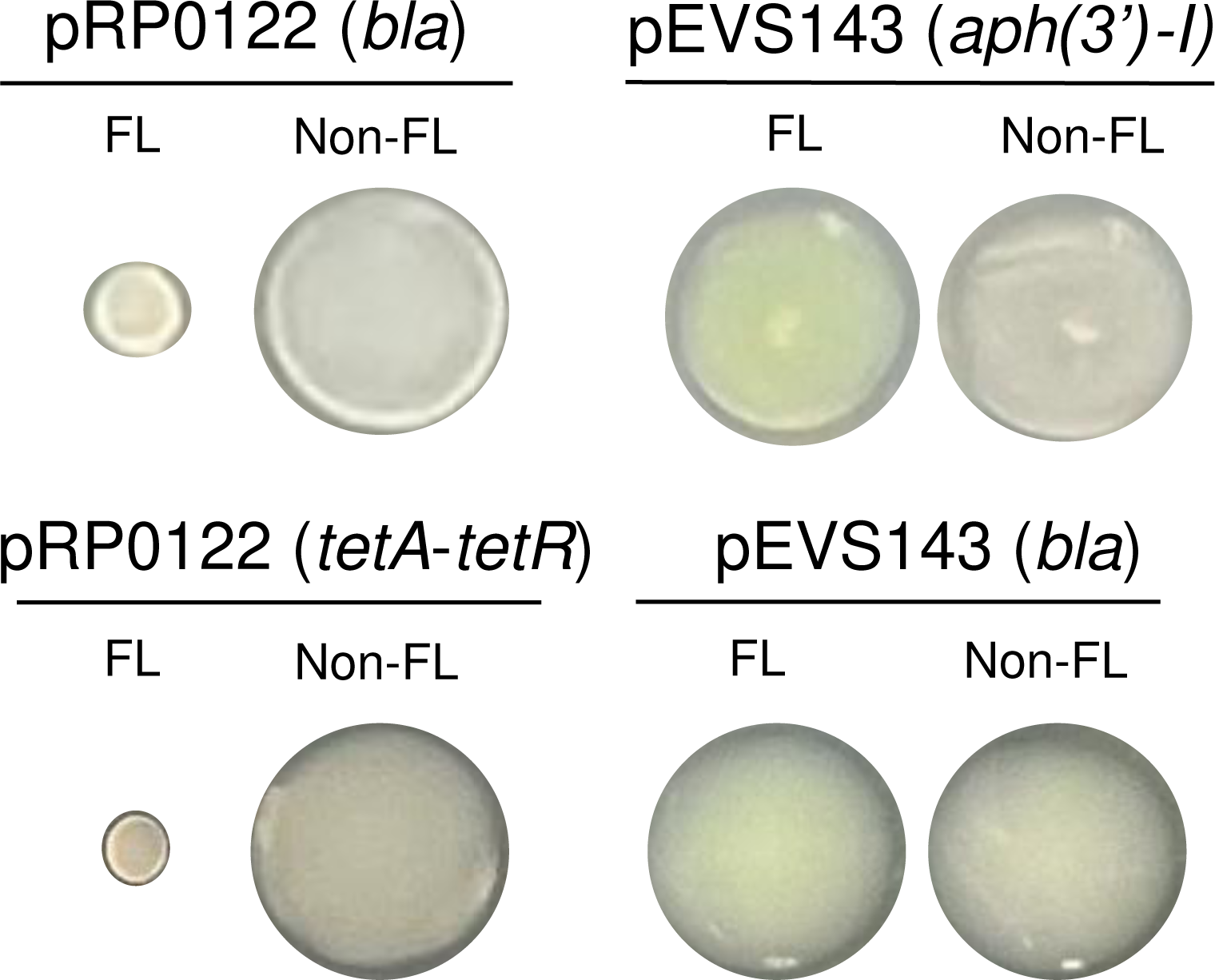
*V. cholerae* WT encoding the indicated plasmid was isolated from a spot colony and individual colonies were grown on non-selective media. Representative images of the visually fluorescent (FL) and non-fluorescent (non-FL) colonies are shown.

Fluorescent and non-fluorescent colonies from the non-selective plates were patch plated on plates containing their cognate antibiotic to measure plasmid maintenance frequency. As expected, fluorescent colonies of the pRP0122 (*bla*), pRP0122 (*tetA-tetR*), pEVS143 (*aph(3’)-I*), and pEVS143 (*bla*) plasmids were able to grow on the selective antibiotic at 100% frequency in both the WT and Δ*vpsL* background (Table 1). Non-fluorescent pEVS143 (*aph(3’)-I*) and pEVS143 (*bla*) colonies were completely unable to grow on selective media, confirming that these colonies had lost their plasmid. While a low frequency of pRP0122 (*bla*) and pRP0122 (*tetA-tetR*) non-fluorescent colonies were able to grow on selective media, the majority of these colonies were unable to grow confirming that plasmid loss had occurred (Table 1). These results confirm that the loss of fluorescence is an accurate estimation for plasmid loss.

**Table 1.** Percent plasmid maintenance by dividing growth on selective media by growth on non-selective media and multiplying by 100.

|  | **Wild Type** |  | **$\Delta vpsL$** |  |
| --- | --- | --- | --- | --- |
|  | ***Percent Growth on Selective Media*** |  | ***Percent Growth on Selective Media*** |  |
| ***Plasmid-Resistance Mechanism*** | ***Fluorescent*** | ***Non-fluorescent*** | ***Fluorescent*** | ***Non-fluorescent*** |
| pRP0122-*bla* | 100% | 10.5% | 100% | 0% |
| pRP0122-*tetA-tetR* | 100% | 8.3% | 100% | 6.9% |
| pEVS143-*aph(3')-I* | 100% | 0% | 100% | 0% |
| pEVS143-*bla* | 100% | 0% | 100% | 0% |

## Discussion

Cooperation is a key element of bacterial survival. Cooperative behaviors, including production of public goods that are available to nearby members, add to the overall success of a community or population. Cooperative behaviors can be either reciprocated (31, 32) or exploited by cheaters (7,8). Cheaters benefit from the production of costly products made by cooperators, and in some cases, outcompete cooperators (13,15). The evolutionary implications of cooperation and cheating are generally considered in the context of ecology and environmental fitness of the species of interest, but such dynamics also exist in laboratory settings and can have significant impacts on experimental outcomes.

In this study, we show that using beta-lactams and the beta-lactamase *bla* to select for plasmid maintenance in a *V. cholerae* colony leads to the evolution of plasmid-free cheaters that were able to thrive in an environment that had been detoxified by cooperative, ampicillin resistant members of the population. As the plasmid we were studying encodes a biosensor for c-di-GMP that can be used to quantify concentrations of this signal in single cells, such plasmid loss has profound implications on interpretation of the experiment. In this case, cells that have lost the plasmid could be mistaken as cells with low intracellular c-di-GMP concentrations.

Based on this finding, we expanded our study to other antibiotic resistance markers, including an aminoglycoside phosphotransferase (*aph(3’)-I*) and a tetracycline antiporter (*tetA-tetR*). Surprisingly, selecting for plasmid maintenance with these additional selection markers and their cognate antibiotic also led to the evolution of plasmid-free cheaters, indicating the potential for plasmid loss during laboratory experiments is not limited to *bla* and ampicillin. Furthermore, the fact that we observed plasmid loss with two plasmid backbones that utilize two distinct origins of replication with different copy numbers demonstrates that laboratory plasmid loss is not specific to only pRP0122 (*bla*).

The emergence of cheater cells on solid agar has recently been demonstrated through the study of persister cells and satellite colonies. Medaney et al showed that *Escherichia coli* harboring a beta-lactamase encoding plasmid secretes the enzyme into the media to create a zone of ampicillin clearance leading to propagation of susceptible, plasmid-free cells in this zone (16). In this study, the authors used a naturally occurring plasmid and concluded that resistant cells were unable to protect sensitive cells unless they were inoculated at high frequency. Our study examined a commonly used laboratory plasmid and found that plasmid-free cells could spontaneously emerge and plasmid containing cells were often present in very low abundance in the colony periphery. Previous studies have demonstrated that beta-lactamase secreting strains could protect ampicillin sensitive strains from antibiotic killing, but these studies explored this dynamic by competing isogenic resistant and sensitive strains (13,15). Importantly, we find that a clonal population of plasmid-encoding cells exhibits plasmid loss in an experimental setting on a rapid time scale during the formation of one colony.

The evolution of cheaters that utilize other antibiotic resistance mechanisms is less studied. While our original hypothesis was that beta-lactam resistance would produce the highest frequency of cheaters based on its extracellular location, we found similar emergence of plasmid-free cells with two additional resistance mechanisms. Tetracycline resistance via antiporter activity showed an equally, and in some cases more severe, ability to permit the growth of cheaters at high frequency. Once tetracycline enters the cell, it chelates metals such as Ca^2+^ and Mg^2+^ and forms a complex (33, 34). This chelation event allows tetracycline to bind to the *tetR* repressor, which frees the operator *tetO* and allows transcription of *tetA*, the antiporter (35, 36). The tetracycline-metal complex is then transported out of the cell. This complex of tetracycline with metals may inhibit re-uptake of tetracycline into the cell through incompatibility with outer membrane protein channels.

We also showed that kanamycin resistance via phosphotransferase activity permitted the growth of cheaters on solid media at a high frequency, implicating phosphotransferase activity as an intracellular public good. Once kanamycin enters the cell, the aminoglycoside(3’)-phosphotransferase enzymatically inactivates the antibiotic. We speculate that this activity depletes active kanamycin in the surrounding growth media, allowing plasmid-free cheaters to emerge. Interestingly, satellite colonies do not emerge when selecting with kanamycin. Since this inactivation occurs in the cell cytoplasm, unlike secreted beta-lactamase, only plasmid-free cells in close proximity benefit from antibiotic inactivation.

Plasmid-free cheaters also arose in a planktonically grown cultures although to a lesser extent than growth on solid surfaces. Cultures in which plasmid maintenance was maintained using tetracycline showed the greatest inhibition of cheaters. Kanamycin selection for plasmid maintenance using the *aph(3’)-I* gene showed a greater inhibition of cheaters in WT *V. cholerae* versus the Δ*vpsL* mutant in planktonic cultures. Further exploration into how this genotype may alter plasmid maintenance is needed, but the major difference between these strains is the ability to form biofilms. Importantly, we found that beta-lactam selection resulted in a high frequency of cheaters during planktonic growth. Various chemostat models have investigated beta-lactamase as a public good that is able to detoxify the liquid environment and permit the growth of cheaters and have detected free beta-lactamase in the growth media (13, 14). Our results add to these findings and demonstrate that plasmid-free cheaters spontaneously arise in a liquid growth medium where beta-lactams are used to maintain plasmids with the *bla* antibiotic selection marker.

Although beta-lactamases represent an extracellular means of antibiotic clearance, we have also demonstrated that two intracellular antibiotic resistant mechanisms promote the evolution of cheater cells in an isogenic population. It is generally assumed that antibiotic selection for plasmids leads to plasmid maintenance in the entire population, but our results suggest this is not always the case. The implications of heterogeneity in experimental bacterial populations could widely impact various genetic experiments such as extrachromosomal complementation, gene expression, or cellular biomarkers. Moreover, as this effect is amplified on solid surfaces, spontaneous plasmid loss may impact experiments on surfaces such as exploring colony morphology (37–39), surface sensing (40–42), motility (43, 44), and biofilm formation studies (45, 46).

One approach to reduce heterogeneity is incorporating cellular biosensors or genetic overexpression constructs into the chromosome using techniques such as Lambda Red recombination (47, 48), multiplex automated genome engineering (MAGE) (49, 50), or multiplex genome editing by natural transformation (MuGENT) (51). Additionally, careful consideration should be taken to reduce the number of generations that liquid cultures are grown in the presence of antibiotic, as significant plasmid loss was measured at 16 hours. Nevertheless, it is important for microbiologists to consider the evolution of plasmid-free cheater cells during plasmid-based experiments and the impacts that such evolution may have on experimental results.

## Materials and Methods

### Strains and Growth Conditions

All strains were grown in Luria Broth (10% tryptone, 5% yeast extract, 10% NaCl) and supplemented with appropriate antibiotics (described below). The wild type of C6706str2 (52) and a Δ*vpsL* (53) knockout in the same background were used for all experiments involving *V. cholerae*. Plasmids were mated into these backgrounds using a laboratory stock of *Escherichia coli* BW29427, a diaminopimelic acid (DAP) auxotroph, harboring pRP0122-Pbe_amcyan_Bc3-4_turborfp, a cellular biosensor encoded on the pRP0122 plasmid used for measuring intracellular c-di-GMP that was received as a gift from Fitnat Yildiz (22, 23), pEVS143 (54), or pEVS143 (*bla*) (pAMB22, construction described below) and pRP0122 (*tetA-tetR*) (pAMB24, construction described below). *E. coli* BW29427 was additionally supplemented with 300 µg/mL of DAP during growth. All cultures were grown at 35°C for 16 hours with shaking at 210 RPM. When strains were grown in the presence of tetracycline, extra care was taken to conceal the growth media from light. For colonies analyzed on solid media, 10 µl of an overnight culture were plated in a single spot and allowed to dry. The plates were incubated statically at 35°C for 16 hours. Colonies grown on solid media were imaged using a Leica MZ6 stereo zoom microscope. For microscopy experiments where pEVS143-*aph(3’)-I* and pEVS143-*bla* were measured using the GFP channel, fluorescence was induced in liquid media by growing the cultures overnight at 35°C for 16 hours in LB supplemented with 60% glycerol, 10% glucose, and 8% lactose. On solid media, these strains were plated on solid LB agar plates containing either 100 μg/mL of ampicillin or kanamycin and 100 µM of isopropyl β-D-1-thiogalactopyranoside (IPTG) and grown statically at 35°C for 16 hours.

### Antibiotic Preparation

Ampicillin (D-(-)-α-Aminobenzylpenicillinsodium salt, Gold Biotechnology), carbenicillin (Carbenicillin (disodium), Gold Biotechnology), and kanamycin (Kanamycin monosulfate, Gold Biotechnology) were prepared at a concentration of 100 mg/mL by dissolving 1g of antibiotic in 10 mL of deionized water. The stocks of these antibiotics were diluted 1:1000 in LB when growing resistant strains for a final working concentration of 100 µg/mL. Tetracycline (Tetracycline hydrochloride, Sigma-Aldrich) was prepared at a stock concentration of 10 mg/mL by dissolving 100 mg of tetracycline in 10 mL of 70% ethanol. The stock of tetracycline was diluted 1:1000 in LB when growing tetracycline resistant strains to achieve a final working concentration of 10 µg/mL.

### Liquid Chromatography Double Tandem Mass Spectrometry

Overnight cultures were grown as described in LB broth containing 100 µg/ml ampicillin. 10 µl of the overnight culture were plated as a spot on solid LB plates containing 100 µg/ml ampicillin and allowed to dry. The plates were incubated at 35°C for 16 hours. A sterile loop was used to collect cells from either the center or edge of the colony and resuspended in LB. Cells were prepared as previously described (55). Briefly, c-di-GMP was extracted using a buffer of 40:40:20 Methanol:Acetonitrile:Water + 0.1N Formic Acid and stored in HPLC water. LC-MS/MS was performed using negative phase electrospray ionization using the Xevo TQ-S located in the MSU RTSF Mass Spectrometry core using settings as previously described (56).

### Solid agar antibiotic challenge

Solid agar with increasing concentrations of ampicillin and carbenicillin were made using LB broth and bacteriologic grade agar (Research Products International, 15 g/L). Stocks of ampicillin and carbenicillin were diluted in the media to achieve final concentrations of 25, 50, 100, 200, and 500 µg/mL. Overnight cultures of each strain were grown from freezer stocks in LB with 100 µg/ml ampicillin at 35°C and 210RPM shaking. 10 µl of overnight culture were plated as a spot onto the agar plates and allowed to dry in a biological safety cabinet. The plates were incubated statically at 35°C for 16 or 24 hours. Spot colonies were imaged using a Leica MZ6 stereo zoom microscope.

### Cloning to Change Antibiotic Resistance Cassettes

Primers (Table S1) synthesized by Integrative DNA Technologies were designed that amplified on either side of the antibiotic resistance cassette that was to be removed, as well as both upstream and downstream from the center of either the origin of replication (p15A *ori*, pEVS plasmids) or *repA* (replication protein A, pRP0122 plasmids). Primers for amplification of the antibiotic resistance marker to be cloned in place of the previous resistance marker were designed with regions of homology to the regions of the plasmid immediately up- and downstream of the previous resistance marker. All products were amplified using Q5-High Fidelity DNA polymerase according to the manufacturer (New England Biolabs). The PCR reactions containing each part of the amplified plasmid backbone were treated with DpnI to remove the methylated parent plasmid. The two backbone halves and the desired antibiotic resistance marker were PCR purified using the DNA Clean & Concentrator kit available from Zymo Research. Purified PCR products were incubated together in a thermocycler with 1X Hi-Fi DNA Assembly master mix (NEB) at 50°C for one hour. The product was transformed into electrocompetent *E. coli* BW29427 cells. Colonies that grew on LB supplemented with DAP (300 µg/mL) and with appropriate antibiotics (100 µg/mL of ampicillin, kanamycin, or 10 µg/mL of tetracycline) were propagated in liquid LB supplemented with DAP and appropriate antibiotics (as described above) and grown at 35°C for 16 hours. The following day, plasmids were prepped from the cell using a Wizard® Plus SV Minipreps DNA Purification kit (Promega). The entire plasmid sequence was obtained using the company Plasmidsaurus. Once verified, the plasmids were conjugated into *V. cholerae* C6706str2 in the wild type and Δ*vpsL* backgrounds for further analyses.

### Fluorescent microscopy

For initial screens of cells plated onto an LB plate containing appropriate antibiotics as a 10 µl spot, cells from the red center and white perimeter of the colony were collected with a sterile loop and resuspended in 1 mL of LB. The cells were normalized to an OD_600_=0.5 in 1X PBS. 5 µl of the resuspension were transferred to a coverslip and placed on a 1% agarose gel pad. Slides were analyzed with a Leica DM5000 using the combination feature to view both brightfield and fluorescent channels. For experiments performed in liquid culture, overnight cultures were diluted to an OD_600_=0.5 in 1X PBS. 5 µl were pipetted onto a cover slide and placed on a 1% agarose gel pad. Cells were captured using SPOT basic software. Post-image analyses were performed using Fiji software.

### Antibiotic Selection Analysis of Plasmid Loss

Strains were grown and 10 µl of culture was plated on antibiotic plates as previously described. After incubation, the 10 µl cell spots were collected and resuspended into 1 mL LB. The suspension was diluted in a 10-fold series and plated onto LB agar. The plates were incubated overnight at 35°C for 16 hours. The following day, isolated colonies were selected and patched onto LB with appropriate antibiotics and an LB plate (no selection) as a control. These plates were incubated as previously described. Plasmid retention frequency was measured based on the total number of patched colonies that grew on the selective plates divided by growth on the non-selective plate and multiplied by 100.

## Acknowledgements

We thank Fitnat Yildiz for the c-di-GMP biosensor plasmid, pRP0122. This work was supported by National Institutes of Health (NIH) grants GM139537 and AI158433 to C.M.W.

